# Sst- and Vip-Cre mouse lines without age-related hearing loss

**DOI:** 10.1101/2024.07.15.603003

**Authors:** Calvin T. Foss, Timothy Olsen, James Bigelow, Andrea R. Hasenstaub

## Abstract

GABAergic interneurons, including somatostatin (SST) and vasoactive intestinal peptide (VIP) positive cells, play a crucial role in cortical circuit processing. Cre recombinase-mediated manipulation of these interneurons is facilitated by commercially available knock-in mouse strains such as Sst-IRES-Cre (Sst-Cre) and Vip-IRES-Cre (Vip-Cre). However, these strains are troublesome for hearing research because they are only available on the C57BL/6 genetic background, which suffer from early onset age-related hearing loss (AHL) due to a mutation of the Cdh23 gene. To overcome this limitation, we backcrossed Sst-Cre and Vip-Cre mice to CBA mice to create normal-hearing offspring with the desired Cre transgenes. We confirmed that in these “CBA Cre” lines, Cre drives appropriate expression of Cre-dependent genes, by crossing CBA Cre mice to Ai14 reporter mice. To assess the hearing capabilities of the CBA Cre mice, we measured auditory brainstem responses (ABRs) using clicks and tones. CBA Cre mice showed significantly lower ABR thresholds compared to C57 control mice at 3, 6, 9, and 12 months. In conclusion, our study successfully generated Sst-Cre and Vip-Cre mouse lines on the CBA background that will be valuable tools for investigating the roles of SST and VIP positive interneurons without the confounding effects of age-related hearing loss.

## 1. Introduction

Cre-Lox recombination tools in mice have enabled cell-type specific manipulations of neuronal activity (reviewed in Luo et al., 2018; Taniguchi et al., 2011; Tsien et al., 1996; Zhang et al., 2011) which have provided important insights into circuit mechanisms in the central auditory system (Bigelow et al., 2019; reviewed in DiGuiseppi and Zuo, 2019; Olsen and Hasenstaub, 2022). These tools have revolutionized our understanding of cortical inhibitory interneurons, including the somatostatin (SST) and vasoactive intestinal peptide (VIP) positive cells, which play crucial roles in cortical function (Bigelow et al., 2019; Natan et al., 2015; Pfeffer et al., 2013; Phillips and Hasenstaub, 2016).

But there is an important limitation: in the majority of commercially available Cre strains, the Cre gene is carried on a C57BL/6 background. C57 mice carry a mutated copy of the Cdh23 gene (Noben-Trauth et al., 2003), which causes age-related hearing loss (AHL; Ison and Allen, 2003; Li and Borg, 1991). This limits the usefulness of these Cre-bearing mice for studies of the auditory system, especially since hearing loss due to the Cdh23 mutation is already present by early adulthood and rapidly worsens over time (Li and Borg, 1991; Willott, 1986). CBA/CaJ background mice do not carry the AHL mutation and have better hearing than C57 mice (Henry and Chole, 1980; Hequembourg and Liberman, 2001; Spongr et al., 1997). The AHL phenotype is recessive, and first generation (F1) crosses of CBA and C57 have normal hearing (Frisina et al., 2011). However, CBA and C57 mice have significant genetic and behavioral differences (Lilue et al., 2018; Sultana et al., 2019), rendering F1 C57-CBA crosses heterozygous in many key alleles. Thus, it is desirable to iteratively backcross AHL-negative C57-CBA hybrids onto the CBA background across multiple generations (>F4) to achieve genetic homogeneity suitable for the multi-generational breeding schemes that are often necessary to leverage the power of Cre (Grove et al., 2016; Markel et al., 1997).

Herein, we describe two Cre knock-in mouse strains which we backcrossed to the CBA/CaJ background to generation F6: Sst^tm2.1(cre)Zjh^/J (“Sst-IRES-Cre”) and Vip^tm1(cre)Zjh^/J (“Vip-IRES-Cre”). We confirmed normal hearing in these strains with auditory brainstem response (ABR) measurements, and normal Sst-Cre and Vip-Cre expression by further crossing these strains with Ai14 Cre-dependent reporter mice. Finally, we made these strains publicly available by donating them to The Jackson Laboratory (JAX).

## 2. Materials and Methods

All procedures were approved by the Institutional Animal Care and Use Committee at the University of California, San Francisco.

### 2.1 Animal Husbandry

We crossed female Sst-IRES-Cre (stock no. 013044) and Vip-IRES-Cre (stock no. 010908) mice with CBA/CaJ (stock no. 000654) male mice, all purchased from JAX. Subsequent generations were genotyped for AHL and Cre, and AHL-negative, Cre-positive females were used to continue the backcrossing. In cases where such females were not available, we instead crossed AHL-negative, Cre-positive males with CBA/CaJ females. We continued backcrosses through generation F6, at which point we donated young males from each strain to JAX for distribution to other investigators as Sst-IRES-Cre (stock no. 037963) and Vip-IRES-Cre (stock no. 037964). We used generation F4 mice to verify normal hearing via ABR measures collected through 12 months of age (described below). We used generation F5 mice to verify Cre expression patterns by crossing male Ai14 mice (Cre-dependent tdTomato; JAX stock no. 007914) to female CBA Cre-bearing mice, and to female C57 Sst-IRES-Cre and Vip-IRES-Cre mice.

Animals were socially housed in cages of 2-5 animals under a 12 h-12 h light-dark cycle. Food and water were provided ad libitum.

### 2.2 Genotyping

Mice were genotyped by Transnetyx (Cordova, TN) for Cdh23 and Sst-Cre or Vip-Cre. Primers for the Cdh23 AHL gene were as follows: forward:

TGCCCTACAGTACTAACATCTACGA, reverse: ACGCAGGACAGGCATTTGT, reporter 1: CTCTCCTCCGGTGAGC, reporter 2: CTCTCCTCCAGTGAGC. Primers for the Sst-IRES-Cre transgene were as follows: forward: GTCAGGTACATGGATCCACTAGTTC, reverse: GCCAGGAGTTAAGGAAGAGATATGG, reporter: CTAGGACAACAATATTGCGGCCG. Primers for the Vip-IRES-Cre transgene were as follows: forward: TCAGGTACATGGATCCACTAGTTCT, reverse: GCACGCTCACCTCTGATTTCA, reporter: AGGCCTCTTCGCGGCCG.

### 2.3 Perfusion and tissue processing

Four CBA Sst-Cre:Ai14, five C57 Sst-Cre:Ai14, five CBA Vip-Cre:Ai14, and eight C57 Vip-Cre:Ai14 mice were euthanized at p36-p41 with sodium pentobarbital (Fatal-Plus, Vortech Pharmaceuticals) and perfused with ice cold, phosphate-buffered saline (PBS) followed by 4% paraformaldehyde (PFA) in PBS. Brains were fixed in 4% PFA overnight and changed to a 30% sucrose solution the following day. After at least one day in sucrose solution, brains were frozen and cut into 50-µm thick coronal slices on a Microm HM 450 sliding microtome. Slices were mounted on slides using Vectashield with DAPI mounting media and imaged using a Keyence BZ-X810 fluorescent microscope. No immunostaining was performed; example images in Figure 1 reflect unamplified tdTomato fluorescence.

**Figure 1.**
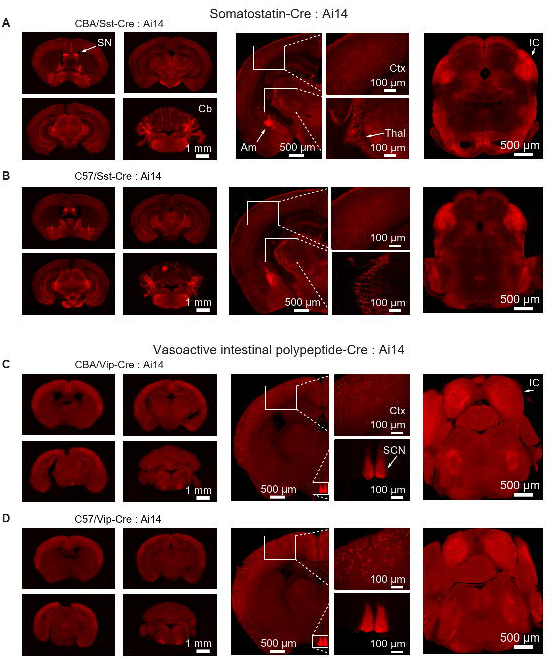
Histological comparison of Cre-driven expression patterns in commercially available and backcrossed Sst-Cre and Vip-Cre mice. A) Example CBA Sst-Cre : Ai14 mouse. i) Low magnification image of coronal brain slices (50 µm thickness) spaced 600 µm apart, in order from rostral to caudal from left to right, top to bottom. ii) Higher magnification image of cortical slice with expanded images of cortex and thalamus showing expression. ii) Higher magnification image showing inferior colliculus expression. B) (i-iii) Example C57 Sst-Cre : Ai14 mouse showing similar expression patterns to A (i-iii). C) Example CBA Vip-Cre : Ai14 mouse. i) Low magnification image of coronal brain slices. ii) Higher magnification image of cortical slice with expanded images of cortex and suprachiasmatic nucleus showing expression. ii) Higher magnification image showing inferior colliculus expression. D) (i-iii) Example C57 Vip-Cre : Ai14 mouse showing similar expression patterns to C (i-iii).

### 2.4 Auditory brainstem response (ABR) measurements

ABR measures were collected at 3, 6, 9, and 12 months of age from eight CBA Sst-Cre mice, nine CBA Vip-Cre mice, and ten C57BL/6J control mice (JAX stock no. 000664). Additional measurements were performed on C57BL/6J mice at 2 months old. Mice were anesthetized with a combination of ketamine (90 mg/kg IP, Ketathesia, HenrySchein) and xylazine (10 mg/kg IP, AnaSed, Akorn) and maintained with supplemental doses of ketamine (25-50 mg/kg IP) as needed. Recordings were conducted inside a sound-attenuation chamber lined with anechoic foam.

Stimuli included clicks (0.1 ms ungated square waves) and pure tone frequencies of 4, 8, 16, 32, and 48 kHz (5 ms sine waves with 0.5-ms cosine^2^ ramps) presented at levels spanning 20–90 dB SPL in 5-dB steps. Each click- and frequency-level combination was repeated 500 times in pseudorandom order with a 30-ms inter-stimulus interval. The stimuli were generated digitally with MATLAB at 192 kHz then converted to analog with a Roland Quad-Capture sound card, amplified by a Tucker-Davis Technologies (TDT) SA1 power amplifier, and delivered through a TDT MF1 speaker in the closed field configuration. Stimulus levels were calibrated with a Brüel & Kjær model 2209 sound level meter and a model 4939 microphone.

Physiological signals were recorded by positioning subdermal silver wire electrodes at the right bulla, vertex (reference), and hindlimb (ground). Continuous voltage traces from the electrodes were amplified by a Medusa preamp and streamed to disk by a RZ2 acquisition system (TDT). The raw signal was bandpass filtered offline (500-3,000 Hz) and averaged across the trials for each stimulus/attenuation pair to obtain the ABR. Example stimuli and ABRs are shown in Figure 2A.

**Figure 2.**
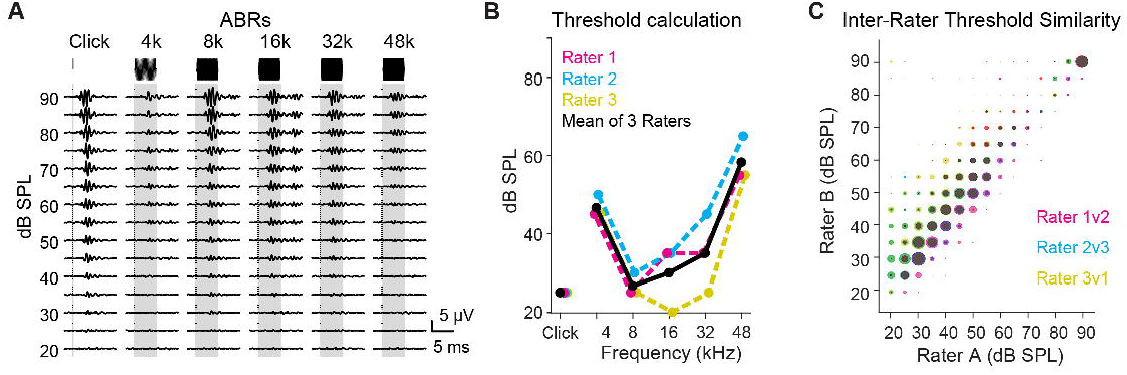
Auditory brain response (ABR) stimuli and analyses. A) Responses to click and pure tone stimuli are averaged to obtain ABR. B) Hearing thresholds are defined as the mean of blinded raters’ assessed thresholds. C) Assessed thresholds for all recordings showed consistency between raters.

### 2.5 Data analysis

Similar to previous studies (e.g., Akil et al. 2016, Ingham et al. 2019), we estimated ABR thresholds by visual inspection of ABR waveforms. Three expert reviewers independently determined the minimum sound level (threshold) at which a recognizable ABR pattern was observed for each recording and stimulus. If no ABR was apparent at any sound level (e.g., high frequency tones in aged C57 mice), threshold was scored as the maximum possible level (90 dB) for purposes of statistical analysis.

Example rater responses are shown in Figure 2. Thresholds across raters were highly correlated with each other as seen in Figure 2D (1v2 r = 0.94, 2v3 r = 0.93, 3v1 r = 0.58; p < 0.0001 for all). We define ABR thresholds as the mean of the three rater values. All threshold judgments were performed blind to animal strain and age.

### 2.6 Statistics

Statistical analyses were performed using MATLAB. Because ABR thresholds are not normally distributed, they were first transformed using the Box-Cox procedure (Box and Cox, 1964). We then used two-way ANOVA to compare thresholds from CBA Sst-Cre and Vip-Cre mice at each age. Thresholds were not significantly different between Sst-Cre and Vip-Cre lines at any age, so the strains were combined for subsequent analyses. We used two-way ANOVA to compare the CBA Cre line thresholds to the C57 thresholds at each age. To control false discovery rate, all p-values were adjusted using the Benjamini-Hochberg procedure (Benjamini and Hochberg, 1995). Following a significant result from the two-way ANOVA, Tukey-Kramer post hoc tests were used to identify significant differences between groups.

## 3. Results

### 3.1 tdTomato expression is similar between CBA Cre:Ai14 and C57 Cre:Ai14 mice

We crossed CBA Sst-Cre and Vip-Cre mice with Ai14 reporter mice to test the function of the Cre modifications in the backcrossed mice. As a comparison, we also crossed C57 Sst-Cre and Vip-Cre mice with Ai14 reporter mice. Coronal sections from CBA Sst-Cre:Ai14 and C57 Sst-Cre:Ai14 mice both show a high concentration of tdTomato+ cells in the septal nucleus (SN), reticular nucleus (Thal), amygdala (Am), inferior colliculus (IC), and cerebellum (Cb), and scattered cells in the cortex (Ctx) (Figure 1 A-B) which parallels expression of Sst-Cre:Ai14 mice from the Allen Mouse Brain Connectivity Atlas (Allen, 2004, a). The CBA Vip-Cre:Ai14 and C57 Vip-Cre:Ai14 slices both show a high concentration of tdTomato+ cells in the suprachiasmatic nucleus (SCN) and inferior colliculus (IC), and scattered cells in the superficial cortex (Ctx) (Figure 1 C-D) which parallels expression of Vip-Cre:Ai14 mice from the Allen Mouse Brain Connectivity Atlas (Allen, 2004, b). Thus, the CBA Sst-Cre and Vip-Cre mouse lines both drive Cre expression in expected brain regions.

### 3.2 CBA Cre mice have normal hearing through 12 months of age

We recorded ABRs using clicks and tones spanning 4-48 kHz (Figure 2) in n = 9 CBA Vip-Cre mice (6 female), n = 8 CBA Sst-Cre mice (2 female), and n = 10 C57 mice (5 female), at 3, 6, 9, and 12 months (Figure 3). We suspected hearing thresholds would be similar between CBA Vip-Cre and Sst-Cre mice. To test this, we performed three-way ANOVA with threshold as the dependent variable, and Cre allele, age, and stimulus type as independent variables. The main effect of strain was not significant (F(1,369) = 1.84, FDR adjusted p = 0.26), as were interactions between strain and age (F(3,369) = 0.58, FDR adjusted p = 0.75), and strain and stimulus (F(5,369) = 0.36, FDR adjusted p = 0.88). Thus, CBA mice bearing these two Cre alleles had similar hearing thresholds and were grouped together for subsequent analyses.

**Figure 3.**
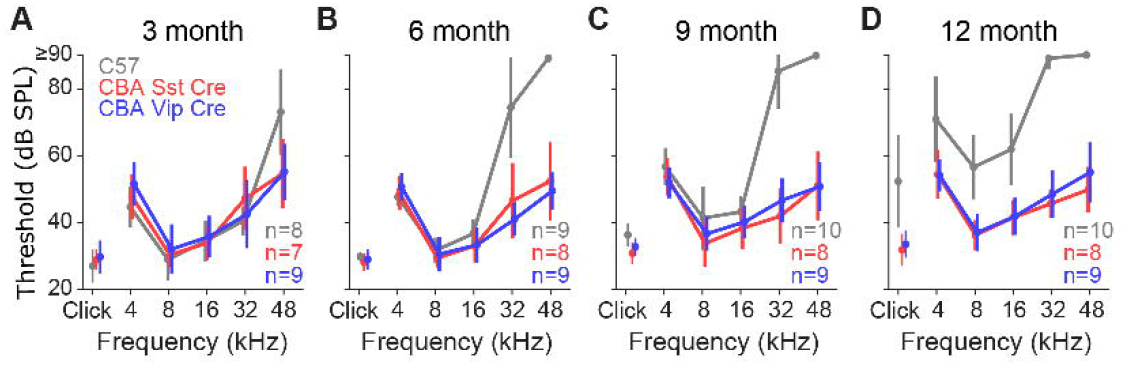
ABR thresholds over time. (A-D) Hearing thresholds for CBA Sst-Cre (red) and CBA Vip-Cre (blue) show consistency over 3, 6, 9, and 12 months while thresholds for C57BL/6J (gray) control mice show elevation over time. Error bars represent one standard deviation.

Differences between CBA Cre and C57 mice were first detectable at 3 months and became progressively more severe with age. At 3 months, the interaction between strain and stimulus was significant (F(5,132) = 3.07, FDR adjusted p < 0.05), consistent with the elevated thresholds seen in C57 mice at 48 kHz (Figure 3A). However, the main effect of strain was not significant (F(1,132) = 3.07, p = 0.4), as were post-hoc tests for any individual stimulus (all p-values > 0.08), suggesting only mild AHL at high frequencies in C57 mice of this age. At all other ages (6, 9, and 12 months), both the main effect of strain and interactions between strain and stimulus were highly significant (all FDR adjusted p-values < 0.001). Post-hoc tests indicated significant differences in hearing thresholds between CBA Cre and C57 mice for 32 and 48 kHz tones at 6 and 9 months (all FDR adjusted p-values < 0.0001), and significant differences across all tone frequencies and clicks by 12 months (all FDR adjusted p-values < 0.001). Together, these outcomes confirmed the severe, progressive AHL phenotype was present in C57 mice but absent in the new CBA Cre strains through at least 1 year of age.

### 3.3 High frequency hearing thresholds in C57 mice are elevated even at 2 months old

Considering early signs of high frequency hearing loss were already evident in C57 mice by 3 months old (Figure 3A), we wondered if such hearing deficits might be detected at an even earlier age. Thus, we collected additional ABR measurements from 2-month-old C57 mice and compared them to the 3-month-old CBA Cre mice (Figure 4). Two-way ANOVA revealed a non-significant effect of strain on thresholds (FDR adjusted p = 0.83) but significant interactions between mouse strain and stimulus (F(5,138) = 2.56, FDR adjusted p < 0.05) indicating a stimulus-dependent effect of strain on hearing. Post-hoc tests revealed significant differences between the groups at 48 kHz (FDR adjusted p < 0.05). Thus, high frequency hearing impairment can be detected in C57 mice as young as 2 months old.

**Figure 4.**
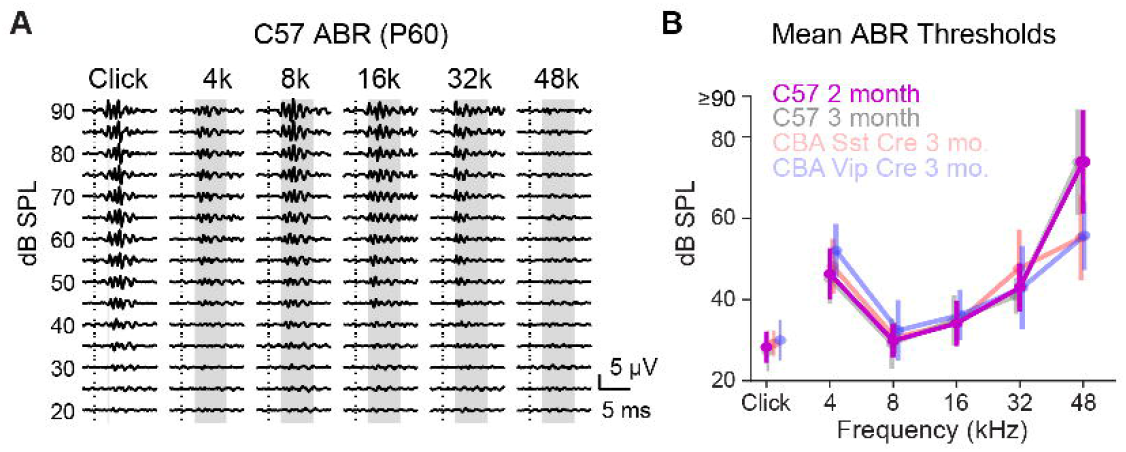
As early as P60, C57-Cre mice show significant hearing deficits compared to older CBAs. A) Representative ABR from a 2 month old C57 mouse shows minimal 48 kHz responses. B) Hearing thresholds for CBA Sst-Cre (light red) and Vip-Cre (light blue) mice at 3 months, and C57BL/6J control mice at 2 months (purple) and 3 months (light gray). Thresholds are elevated for high frequencies in C57 mice at 2 and 3 months. Error bars represent one standard deviation.

## 4. Discussion

Sst-Cre and Vip-Cre transgenic mouse lines on C57 backgrounds have provided numerous insights into cortical microcircuit structure and function by enabling targeted manipulation of VIP- and SST-positive interneurons. However, the Cdh23 mutation carried by these strains confers an AHL phenotype extensively documented by previous studies (Burghard et al., 2019; Frisina et al., 2011; Henry and Chole, 1980; Henry and Lepkowski, 1978; Hequembourg and Liberman, 2001; Ison and Allen, 2003; Kane et al., 2012; Li and Borg, 1991; Mikaelian et al., 1974; White et al., 2000; Willott, 1986). Our findings confirm previous work suggesting AHL is first detectable at high frequencies by 2-3 months old, and progressively worsens with age such that thresholds are severely elevated at high frequencies by 6 months old, and across all frequencies by 12 months old. This early-onset, progressive AHL phenotype hinders the utility of C57-background strains for studies of the auditory pathway and related structures, especially for studies of aging-related changes in these structures.

To circumvent these limitations, we generated novel Sst-Cre and Vip-Cre lines by backcrossing onto a CBA background that does not carry the Cdh23 mutation. We verified the absence of AHL in the new strains by conducting longitudinal ABR recordings through 12 months of age (Figures 3-4). We also histologically confirmed the expected expression patterns of Cre-dependent genes via crosses with Ai14 reporter mice (Figure 1). The new strains thus provide new tools for studying circuit mechanisms of auditory processing and multisensory integration across the lifespan without potential confounds due to progressive AHL. Similar approaches were used by Beebe et al. (2020) and Lyngholm and Sakata (2019) to generate ChAT-Cre and Cre-dependent channelrhodopsin2 strains without AHL.

A minor caveat regarding the use of CBA mice for hearing studies is that males may develop mild adult onset diabetes-obesity syndrome, which has been associated with elevated thresholds in several previous studies (Fujita et al., 2015; Hwang et al., 2013; Lee et al., 2008; Pålbrink et al., 2020). These influences tend to be modest, especially in comparison to AHL caused by Cdh23 mutation in C57 mice. Nevertheless, researchers specifically focusing on sex differences in aged CBA mice may wish to account for potential influences of metabolic status on hearing.

In summary, we produced new Sst-Cre and Vip-Cre mouse strains on CBA backgrounds, then confirmed experimentally they lack AHL and feature the expected

Cre expression patterns. These mice are now publicly available from JAX as Sst-IRES-Cre (stock no. 037963) and Vip-IRES-Cre (stock no. 037964).

## Funding

This work was supported by the National Institutes of Health Grants R01NS116598 and R01MH122478 (to A.R.H.), Hearing Research Inc. (to A.R.H.), the Coleman Memorial Fund (to A.R.H.), and NIA R01AG078132 (to J.B.).

## Declaration of competing interest

The authors declare no competing interests.

## CRediT authorship contribution statement

**Calvin T. Foss:** Methodology, Formal analysis, Investigation, Writing - Original Draft, Writing - Review & Editing, Visualization. **Timothy Olsen:** Writing - Review & Editing, Software, Validation, Formal analysis. **James Bigelow:** Writing - Review & Editing, Software, Validation, Formal analysis, Funding acquisition. **Andrea R. Hasenstaub:** Conceptualization, Methodology, Writing - Review & Editing, Software, Supervision, Project administration, Funding acquisition.

## Acknowledgements

We thank Natalia Santiago and Drs. Christoph Schreiner, Congcong Hu, Philine Marchetta, Toshiaki Suzuki, and Benjamin Rakela for their support with data collection, analysis, and writing.

## Notes

### Competing Interest Statement

The authors have declared no competing interest.

